# DiffGR: Detecting Differentially Interacting Genomic Regions from Hi-C Contact Maps

**DOI:** 10.1101/2020.08.29.273698

**Authors:** Huiling Liu, Wenxiu Ma

## Abstract

Recent advances in Hi-C techniques have allowed us to map genome-wide chromatin interactions and uncover higher-order chromatin structures, thereby shedding light on the principles of genome architecture and functions. However, statistical methods for detecting changes in large-scale chromatin organization such as topologically-associating domains (TADs) are still lacking. We proposed a new statistical method, DiffGR, for detecting differentially interacting genomic regions at the TAD level between Hi-C contact maps. We utilized the stratum-adjusted correlation coefficient to measure similarity of local TAD regions. We then developed a non-parametric approach to identify statistically significant changes of genomic interacting regions. Through simulation studies, we demonstrated that DiffGR can robustly and effectively discover differential genomic regions under various conditions. Furthermore, we successfully revealed cell type-specific changes in genomic interacting regions in both human and mouse Hi-C datasets, and illustrated that DiffGR yielded consistent and advantageous results compared with state-of-the-art differential TAD detection methods. The DiffGR R code is published under the GNU GPL ≥ 2 license and is publicly available at https://github.com/wmalab/DiffGR.

## 1 Introduction

Recent developments of chromatin conformation capture (3C)-based techniques—including 4C [1], 5C [2], Hi-C [3–5], ChIA-PET [6], and Hi-ChIP [7]—have allowed high-throughput characterization of pairwise chromatin interactions in the cell nucleus, and provided an unprecedented opportunity to investigate the three-dimensional (3D) chromatin structures and to elucidate their roles in nuclear organization and gene expression regulation. Among these techniques, Hi-C and its variants [8–10] are of particular interest because of their ability to map chromatin interactions at a genome-wide scale.

A Hi-C experiment yields a symmetric contact matrix in which each entry represents the chromatin contact frequency between the corresponding pair of genomic loci. A particularly important characteristic of Hi-C contact matrices is the presence of the topologically-associating domains (TADs), which are functional units of chromatin with higher tendency of intra-domain interactions [11]. TADs are largely conserved across cell types and species. Moreover, CTCF and other chromatin binding proteins are enriched at the TAD boundaries, indicating that TAD boundary regions form chromatin loops and play an essential role in gene expression regulation [11, 12].

Several computational methods have been developed to detect TADs in Hi-C contact maps. These methods can be categorized into two groups: one-dimensional (1D) statistic-based methods and two-dimensional (2D) contact matrix-based methods [13]. Of these, 1D statistic-based methods often take a sliding window approach along the diagonal of Hi-C contact matrix and compute a 1D statistic for each diagonal bin to detect TADs and/or TAD boundaries. For instance, Dixon et al. [11] introduced a statistic named directionality index (DI) to quantify whether a genomic locus preferentially interacts with upstream or downstream loci and developed a hidden Markov model to call TADs from DIs. Later, Crane et al. [14] proposed a novel TAD detection method, which computes an insulation score (IS) for each genomic bin by aggregating chromatin interactions within a square sliding through the diagonal and then searches for the minima along the IS profile as TAD boundaries. Unlike the 1D statistic-based methods which calculate statistics using local information, the 2D contact matrix-based methods utilize global information on the contact matrix to capture TAD structures. For example, the Armatus algorithm [15] identifies consistent TAD patterns across different resolutions by maximizing a quality scoring function of domain partition using dynamic programming. In addition, Lévy-Leduc et al. [16] proposed a TAD boundary detection method named HiCseg, which performs a 2D block-wise segmentation via a maximum likelihood approach to partition each chromosome into its constituent TADs. Later, Wang et al. [17] introduced a clustering-based TAD calling method CHDF, which optimized the clusters of the contact matrix by dynamic programming with an objective function combining the sum of squared error and a penalty term in favor to domain regions with higher frequency of interactions. Recently, several review papers have quantitatively compared the performances of the aforementioned TAD-calling methods and demonstrated that HiCseg detects a stable number of TADs against changes of sequencing coverage and maintains the highest reproducibility among Hi-C replicates across all resolutions when compared with other TAD-calling methods [18–20].

With the fast accumulation of Hi-C datasets, there has been a growing interest in performing differential analysis of Hi-C contact matrices. To date, several computational tools have been developed for comparative Hi-C analysis, but the majority of them focused on the identification of differential chromatin interactions (DCIs), which represent different chromatin looping events between two Hi-C contact maps. In early studies, the most common strategy for DCI detection was to use the fold change values between two Hi-C contact maps. For instance, Wang et al. [21] used a simple fold-change strategy to detect the influence of estrogen treatment on chromatin interactions in MCF-7 Hi-C samples. Additionally, Dixon et al. [22] utilized the fold change values of chromatin interactions to train a random forest model to discover the epigenetic signals that were more predictive of changes in interaction frequencies. In addition to these fold change-based approaches, another commonly utilized method for detecting DCIs was the binomial model implemented by the HOMER software [23]. In contrast, in more recent studies, count-based statistical methods, such as edgeR [24] and DESeq [25], have been adopted to identify pairwise chromatin interactions that show significant changes in contact frequencies. Among them, Lun and Smyth [26] presented a tool named diffHic for rigorous detection of differential interactions by leveraging the generalized linear model (negative binomial regression) of edgeR, and demonstrated that edgeR outperformed the binomial model. Later, Stansfield et al. [27] introduced MD normalization and performed Z-tests to detect statistically significant DCIs. While all these methods assumed independence among pairwise interactions, which holds true only in coarse-resolution Hi-C maps, Djekidel et al. [28] presented a novel method, named FIND, that takes into account the dependency of adjacent loci at finer resolutions. Briefly, FIND utilizes a spatial Poisson process model to detect DCIs that show significant changes in interaction frequencies of both themselves and their neighborhood bins. Lastly, Cook et al. [29] introduced ACCOST to identify differential chromatin contacts by extending the DESeq model used in RNA-seq analysis and repurposing the “size factor” to account for the notable genomic-distance effect in Hi-C contact matrices.

In the cell nucleus, chromatin is organized at multiple levels, ranging from active and inactive chromosomal compartments and sub-compartments (on a multi-Mb scale) [3, 9], TADs (0.5–2 Mb on average) [11], to fine-scale chromatin interacting loops [8, 9]. Chromatin structures also exhibit multi-scale differences among different cell types in their compartments, TADs, and chromatin loops. Among these, changes in TAD organizations are of particular interest as TADs are strongly linked to cell type-specific gene expression [11]. For example, Taberlay et al. [30] have shown that genomic rearrangements in cancer cells are partly guided by changes in higher-order chromatin structures, such as TADs. They discovered that some large TADs in normal cells are further segmented into several smaller TADs in cancer cells, and these changes are tightly correlated with oncogene expression levels. Current differential analyses of TAD structures between different cell types and conditions are limited to the detection of TAD boundary changes. Recently, Chen et al. [13] proposed a TAD boundary detection approach named HiCDB, which is constructed based on local measures of relative insulation and multi-scale aggregation. In addition to calling TAD boundaries in single Hi-C sample, HiCDB also provides differential TAD boundary detection using the average values of relative insulation across multiple samples. Later, Cresswell and Dozmorov [31] developed TADCompare, which uses a spectral clustering-derived metric named eigenvector gap to identify differential and consensus TAD boundaries and track TAD boundary changes over time., TADreg [32] introduced a versatile regression framework which generalizes the insulation score by estimating the relative insulating effects of genomic loci and adding a sparsity constraint. The TADreg framework was designed for TAD boundary detection, but also allowed differential TAD analysis across various conditions. The HiCDB, TADCompare and TADreg methods focused on detecting changes in TAD boundaries rather than changes in chromatin organization within TADs. However, differential TAD boundaries do not necessarily indicate differential chromatin conformation within those regions. First, Hi-C contact matrices are often sparse and noisy, which might lead to unstable detection of TAD boundaries. Second, chromatin interactions within a TAD could be strengthened or weakened in another Hi-C sample, which would suggest different patterns of chromatin organization within the same TAD region. Unfortunately, few methods have been developed to detect differential TAD regions instead of boundaries. Recently, the Hi-C pre-processing and analysis tool HiCExplorer [33–35] expanded its functions to capture differential TAD regions by comparing the precomputed TAD regions on the target Hi-C map with the same regions on the control map by accounting for the information in both intra-TAD and inter-TAD regions. However, such comparison was only limited to the precomputed genomic regions in only one of the Hi-C conditions. Thus, appropriate statistical methods for detecting differentially interacting regions by considering TAD regions across both conditions are still lacking.

To tackle this problem, we developed a novel statistical method, DiffGR, for detecting differential genomic regions at TAD level between two Hi-C contact maps. Briefly, DiffGR utilizes the stratum-adjusted correlation coefficient (SCC), which effectively eliminates the genomic-distance effect in Hi-C data, to measure the similarity of local genomic regions between two contact matrices. Subsequently, DiffGR applies a nonparametric permutation test on those SCC values to detect genomic regions with statistically significant differential interactions. We demonstrated, through simulation studies and real data analyses, that DiffGR can effectively and robustly identify differentially interacting genomic regions at TAD level.

## 2 Methods

The DiffGR method detects differentially interacting genomic regions in three steps, as shown in Figure 2A and described below in Sections 2.1-2.3. In addition, the simulation settings are outlined in Section 2.4 and real data preprocessing and analyses are described in Section 2.5.

**Figure 1:**
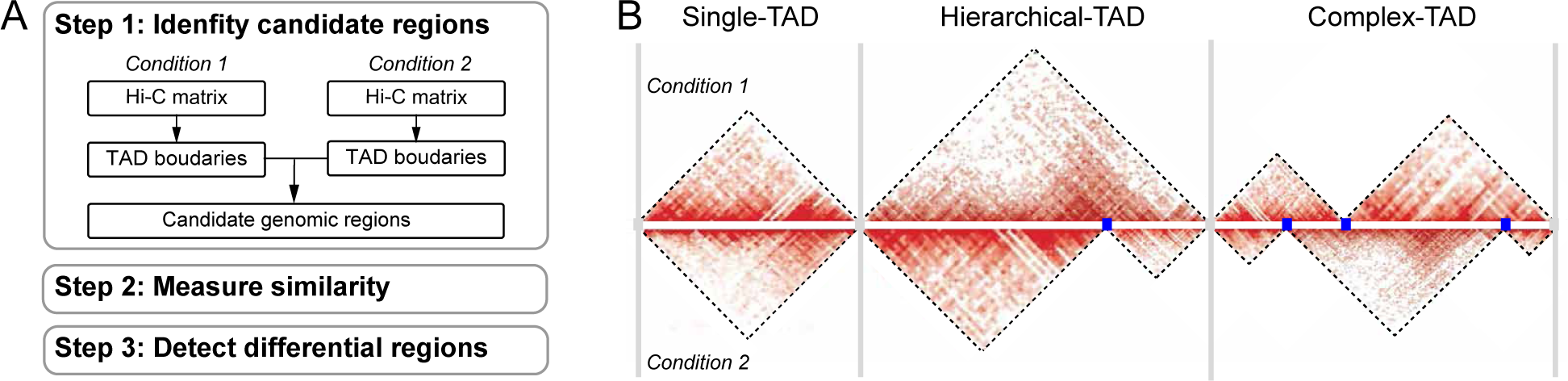
Overview of DiffGR. A. Workflow of the DiffGR algorithm. B. Illustration of three candidate types of differential genomic regions. The gray vertical bars represent the common TAD boundaries between two conditions, which partition the genome into three types of candidate regions. The blue points stand for unique TAD boundaries in only one of the two conditions.

**Figure 2:**
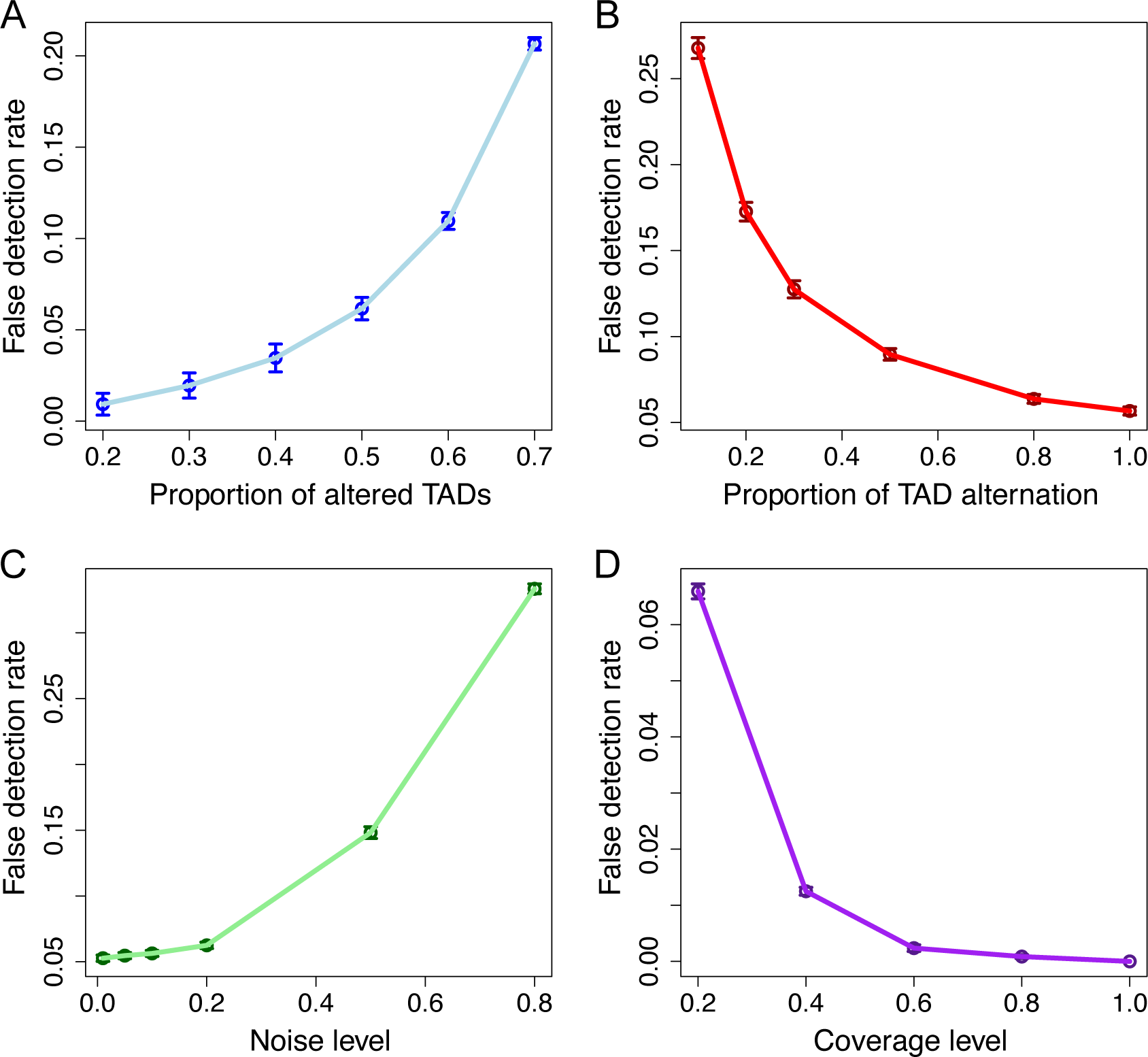
Performance of single-TAD simulations. The curves display the mean false detection rates at different levels of A. proportion of altered TADs, B. proportion of TAD alternation, C. noise, and D. sequencing coverage. Vertical bars represent 95% confidence intervals.

### 2.1 Identifying candidate genomic regions

Suppose we have two sets of Hi-C data and their corresponding contact frequency matrices as the input. First, we detect the TAD boundaries in each Hi-C data, separately. Specifically, we apply HiCseg [16] to the raw contact matrices and obtain the corresponding TAD boundaries. Note that in this step one can change HiCseg with any other credible TAD caller, such as CDHF [17] or TADreg [32], whose detected TADs satisfy the non-overlapping and continuous properties. We choose HiCseg because it has been shown that HiCseg produces more robust and reliable TAD boundaries than other TAD-calling methods [18, 20, 32]. We next combine the TAD boundaries from both Hi-C contact maps to identify the candidate genomic regions for subsequent analyses. TAD boundaries within two-bin distance are considered to be a common boundary shared by both Hi-C datasets and replaced by the middle bin locus. We then partition the genome into non-overlapping candidate regions using the common TAD boundaries, and categorize these candidate regions into the following three groups: (1) single-TAD candidate regions, (2) hierarchical-TAD candidate regions, and (3) complex-TAD candidate regions, as illustrated in Figure 2B.

We expected different patterns of differential features in these three kinds of candidate genomic regions. As to the differential single-TAD region, we would expect strength changes occurred in such areas. For differential hierarchical-TAD regions, one large interacting domain could be evidently split into two or more sub-domains, or vice versa, boundaries between TADs disappeared and thus the corresponding domains merged in one of the contact maps. Lastly, domains might be split, merged, or shifted in a more complicated manner thereby constructing an entirely new structure, which would be defined as differential complex-TAD regions. Unlike differential single-TAD regions, the differential hierarchical-TAD and complex-TAD regions represent more disruptive changes in the 3D structure of the chromatin.

### 2.2 Measuring similarity of candidate regions between two Hi-C contact maps

In the second step, we evaluate the similarity of each candidate region between the two samples. Suppose a candidate genomic region is bounded by two common TAD boundaries shared by both Hi-C maps, and contains *k* unique TAD boundaries in either one of the two Hi-C maps (shown as blue points in Figure 2B). In the single-TAD candidate region, *k* = 0; in the hierarchical-TAD or complex-TAD candidate regions, *k >*= 1. For each candidate region, we consider all 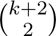 possible (sub)TADs, which are separated by any pair of TAD boundaries within that region, as potential differential TADs. For each potential differential TAD, we calculate the stratum-adjusted correlation coefficient (SCC) [36] rather than the standard Pearson or Spearman correlation coefficients (CCs) to measure the similarity of intra-TAD chromatin interactions between two Hi-C samples. The advantages of using SCC instead of standard CCs are shown in Supplementary Results S1.

The SCC metric was introduced by Yang et al. [36] as a measure of similarity and reproducibility between two Hi-C contact matrices. To account for the pronounced distance-dependence effect in Hi-C contact maps, chromatin contacts are first stratified into *K* stratum according to the genomic distances of the contacting loci pairs, and the correlation coefficients of contacts within each stratum are calculated between two samples. These stratum-specific correlation coefficients are then aggregated to compute the SCC value using a weighted average approach, where the weights are derived from the Cochran-Mantel-Haenszel (CMH) statistic [37]. That is, the SCC *ρ* is calculated as

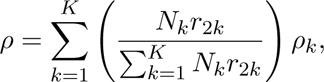

where *N_k_*is the number of elements in the *k*-th stratum, *r*_2*k*_ is the product of standard deviations of the elements in the *k*-th stratum of both samples, and *ρ_k_* denotes the correlation coefficient of the *k*-th stratum between two samples.

The original SCC metric is computed using the intra-chromosomal contact matrices with a predefined genomic distance limit. The resulting value has a range of [*−*1, 1] and can be interpreted in a way similar to the standard correlation coefficient. Here we use SCC as a local similarity measurement to evaluate each potential differential TAD between two Hi-C samples. In the SCC calculation, an upper limit of genomic distance is set to 10 Mb because TADs are commonly smaller than 10 Mb and distal interactions over a genomic distance larger than 10 Mb are often sparse and highly stochastic. In addition, as the sparsity of Hi-C matrices might affect the precision of SCC values, the loci pairs with zero contact frequencies in both samples are excluded from the calculation.

Hi-C contact maps are often sparse due to sequencing coverage limits and contain various systematic biases. To solve these issues, when preprocessing the Hi-C contact matrices, we first smooth each contact map by a 2D mean filter [36], which substitutes the contact count observed between each bin pair by the average contact count in its neighborhood. This smoothing process improves the contiguity of the TAD regions with elevated contact frequencies, thereby enhancing the domain structures. Next, we utilize the Knight-Ruiz (KR) normalization [38] on the smoothed matrices to remove potential biases.

### 2.3 Detecting statistically significant differential regions

In the third step, we identify differential genomic regions by first finding differential TADs within these candidate regions. In each candidate genomic region, we calculate the SCC values for all potential differential TADs as described above. Then we develop a nonparametric permutation test to estimate the *p*-values for these local SCC values. Additionally, we propose a quantile regression strategy to speed up the permutation test (see details in Supplementary Method in File S1). Finally, we consider a candidate region to be a differentially interacting genomic region, if at least one TAD within that region exhibits a statistically significant difference between the two samples and the size of the largest differential TAD meeting this criterion is greater than one third of the length of the entire candidate region. The longest differential TADs within the detected differentially interacting genomic regions are defined as the noticeable differential areas.

Specifically, we perform the following nonparametric permutation test for each unique TAD size, as the local SCC values are calculated for all potential differential TADs of various sizes.

Suppose *s* is a potential differential TAD whose length is *l_s_* and SCC value between two Hi-C samples is *ρ_s_*. To assess the statistical significance of the observed SCC value *ρ_s_*, the null distribution of SCC values for TADs of the same size is estimated via the following permutation procedure. To generate a random TAD with length *l_s_*, we first randomly select *l_s_* positions from main diagonal of Hi-C contact matrix, then *l_s_ −* 1 position from the first off-diagonal, …, and lastly 1 position from the (*l_s_ −* 1)-th off-diagonal. We subsequently extract contact counts of these randomly selected positions from the two Hi-C contact matrices to construct the permuted TAD pair and calculate its SCC value. We repeat the above random TAD generation step *N* times (*N* = 2000) and obtain the corresponding SCC values 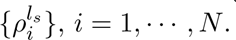 Then the *p*-value of the observed SCC value *ρ_s_* can be computed as:

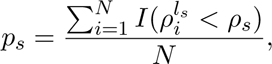

where *I*(*·*) is the indicator function. Lastly, we compare the *p*-values with a pre-defined significance level *α* (by default *α* = 0.05) to determine differential TADs meeting the significance threshold. Note that the permutation framework accounts for the multiple testing correction using the Benjamini-Hochberg procedure [39].

One potential issue of this permutation framework is the false detection of significantly differential TADs when the two samples are highly similar (e.g., biological replicates from same experiment). This is because the high similarity between biological replicates would lead to high SCC values of the corresponding random TAD patterns. As a result, some non-differential TADs with relatively low SCC values would be falsely detected as differential ones. In order to reduce the number of false positives, we provide an option to filter the p-values *p_s_* by an empirical or automatically calculated threshold. This optional filtering step allows us to pre-specify the meaningful SCC between the two Hi-C datasets that should be reached in order to call a differential TAD truly significant.

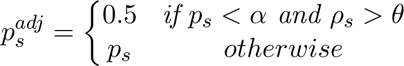

The threshold *θ* can normally be defined as 0.85, which corresponds to a clear margin separating non-replicates from biological/pseudo-replicates in the whole-chromosome similarity comparison between multiple cell lines [40]. Alternatively, *θ* can be calculated automatically as 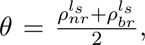 where 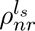 represents the mean *α* quantile of SCCs between non-replicate data and 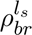 is the mean *α* quantile of SCCs between their corresponding biological/pseudo-replicate data. Here, we call matrices from different cell lines as non-replicates, matrices from the same cell type as biological replicates, and matrices sampled from pooled biological replicates as pseudo-replicates.

### 2.4 Simulation settings

To evaluate the performance of the DiffGR method, we conducted a series of simulation experiments by varying the proportion of altered TADs, proportion of TAD alternation, noise level, and sequencing coverage level. Specifically, we utilized the published chromosome 1 contact matrix of K562 cells at 50-kb resolution [9] as the original Hi-C data and simulated the altered Hi-C contact matrices as described below.

#### 2.4.1 Single-TAD alternation

Since TADs are conserved genomic patterns and TAD boundaries are relatively stable across cell types and even across species [11], our simulations primarily focused on the scenarios of single-TAD alternations. Suppose we had an original Hi-C contact matrix *M* and its identified TAD boundaries. Each of our simulated Hi-C matrices contained two components: the signal matrix *S* and the noise matrix *N*, with a certain signal-to-noise ratio.

First, to construct the signal matrix *S*, we randomly selected a subset of TADs from original contact matrix to serve as the true differential TADs. Then we replaced a certain portion of contact counts in each selected TAD by randomly sampling contact counts from the corresponding diagonals of the contact matrix. Second, we simulated the noise matrix *N* which represents the random ligation events in Hi-C experiments. Briefly, we generated these contacts by randomly choosing two bins, *i* and *j*, and adding one to the entry *N_ij_* in the noise matrix. The probability of sampling each bin in the bin pair was set proportional to the marginal count of that bin in the original matrix. The sampling process was repeated *C* times, where *C* was the total number of contacts in the original Hi-C contact matrix *M*. The resulting random ligation noise matrix *N* contained the same number of contacts as the original contact matrix *M*.

To summarize, we had the following parameters in our single-TAD simulations.

- proportion of altered TADs. Using HiCseg, we detected 189 TADs with a mean size of 1.2 Mb in the original K562 chromosome 1 contact matrix (Supplementary Figure S1). By default, we set the proportion of altered TADs to be 50%, which can vary from 20% to 70%.
- proportion of TAD alternation. In the default setting, we substituted all contact counts in the selected TADs by random counts permuted from the matching diagonals in Hi-C maps. To reduce the degree of intra-TAD alternation, we gradually decreased the proportion of randomly substituted intra-TAD contacts from 100% to 10%.
- noise level, i.e., the ratio between the noise and signal matrices. The noise level was set to 10% by default, and varied from 1% to 80%.

For each simulation parameter setting, we generated 100 altered Hi-C contact matrices to compare against the original contact matrix. To evaluate the accuracy of the detection results, we used the false detection rate which defines as inaccurate percentage and is computed as 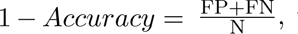 where FP denotes the falsely detected differential regions, FN represents the the falsely detected non-differential regions, and N is the total number of candidate regions being tested.

#### 2.4.2 Hierarchical-TAD alternation

In addition to single-TAD alternation, we also simulated the alternation pattern of hierarchical TADs. We randomly selected 50% of the large TADs whose size was greater than 10 bins in the signal matrix to serve as the true differential TADs. For each of the selected large TAD, we chose a random subTAD boundary to split it into two smaller subTADs (each with size *>* 5 bins). We then replaced all inter-subTAD contact counts by randomly sampled counts in Hi-C maps. Next, we validated the performance of DiffGR under the hierarchical-TAD condition with respect to different noise levels similar to the single-TAD simulations. Because the complex-TAD condition has complicated TAD boundaries between two samples and occurs less frequently in real data, we did not generate simulation data for this condition.

#### 2.4.3 Simulating low-coverage contact matrices

Low sequencing depth of Hi-C experiments would lead to low-coverage and sparse contact matrices, thus it could potentially affect the performance of the detection of differentially interacting regions. To simulate low-coverage contact matrices, we started with a deep-sequenced Hi-C contact map obtained from human GM12878 cells [9], and down-sampled the contact counts to generate lower-coverage matrices. Specifically, for each non-zero contact count *M_ij_* in the original matrix, we assumed that the simulated contact count follows a binomial distribution 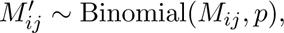 where the binomial parameter *p* = {0.2, 0.4, 0.6, 0.8, 1.0} represents the relative coverage level of the down-sampled contact matrix *M’*. In addition, 10% noise were added to the down-sampled matrices.

### 2.5 Real data preprocessing steps

In our real data analysis, we used two published Hi-C datasets by Rao et al. [9] (GEO accession GSE63525) and Dixon et al. [11](GEO accession GSE35156). The Rao et al. [9] dataset include five human cell types: B-lymphoblastoid cells (GM12878), mammary epithelial cells (HMEC), umbilical vein endothelial cells (HUVEC), erythrocytic leukemia cells (K562), and epidermal keratinocytes (NHEK). The GM12878 dataset contains two replicates, which were also pooled together in cell type-specific comparison. The Dixon et al. [11] dataset are from mouse embryonic stem (ES) and cortex cells. Two replicates from mouse ES cells were merged together in cell type-specific comparison. We applied DiffGR to detect differential genomic regions between each pair of cell types at 25-kb, 50-kb, and 100-kb resolutions. Since some of these Hi-C datasets were not deeply sequenced, the local variations introduced by low sequencing coverage made it challenging to capture large domain structures, especially in fine-resolution analyses. Therefore, to enhance the domain structures, all contact matrices were first preprocessed by a 2D mean filter smoothing and then normalized by the KR method to eliminate potential biases.

In addition to Hi-C contact maps, ChIP-seq and RNA-seq data from the same cell lines were also included in real data analyses. For ChIP-seq analysis, CTCF and histone modification (H3K4me2, H3K9me3, H3K27ac, and H3K27me3) datasets from five human cell lines in Rao et al. [9] were obtained from the ENCODE project [41, 42] (https://www.encodeproject.org/). The ChIP-seq files were in BAM format. The ChIP-seq peaks were called by MACS2 [43] and stored as narrowpeak/broadpeak BED format for the subsequent analyses. In addition, RNA-seq datasets were also obtained from the ENCODE project [42] for human GM12878 and K562 cells (GEO accession GSE78552 and GSE78625) in read count format, and for mouse ES and cortex cells (GEO accession GSM723776 and GSM723769) in FPKM format.

## 3 Results

### 3.1 DiffGR accurately detected single-TAD differences in simulated datasets

To validate the accuracy and efficiency of our DiffGR method, we first generated pairs of original and simulated Hi-C contact matrices, where a given proportion of TADs in the simulated contact matrices were altered (see Methods). We used the intra-chromosomal contact matrix of chromosome 1 in K562 cells at 50-kb resolution to serve as the original contact matrix. At the default setting, we altered 50% of the original TADs by completely replacing the intra-TAD contact counts by randomly sampled counts outside the TAD regions. In addition, we added 10% random-ligation noise into the altered contact matrices.

We first simulated Hi-C matrices with various proportions of altered TADs (20%, 30%, 40%, 50%, 60%, and 70%). With each proportion setting, we completely mutated the intra-TAD counts and added 10% noise, and repeated this simulation procedure 100 times. As expected, the performance of the DiffGR method depended on the proportion of altered TADs. As shown in Figure 2A and Supplementary Table S1, when the proportion of altered TADs changed from 20% to 70%, the false detection rate increased from 0.01 to 0.21. One possible explanation of this observed trend is that when the majority of TADs were altered, the large differences between the original and altered matrices would affect the permutation test and therefore lead to inaccurate detection. However, differential TADs rarely exist in large proportion in real data. The false detection rates of our method remained below 0.07 when the proportion of altered TADs was smaller than or equal to 50%, which demonstrated that our method can accurately and reliably detect single-TAD differences under these conditions.

In the default simulation setting, we completely altered the selected TADs by substituting all intra-TAD contact counts by randomly sampled counts from the matching diagonals outside the TADs. To investigate the influence of the degree of TAD alternation on the DiffGR performance, we generated a series of simulated contact matrices, in which half of original TADs were altered and the proportion of intra-TAD alternation varied from 10%, to 20%, 30%, 50%, 80%, and 100%. In theory, TADs with higher degrees of alternation are easier to identify, whereas TADs with minor changes remain difficult to be detected. As illustrated in Figure 2B and Supplementary Table S2, the performance of DiffGR improved resulting in higher accuracy as the percentage of randomly substituted counts in altered TADs increased. Even with the most challenging case where only 10% of the intra-TAD counts were altered, the accuracy of our method was 0.73, suggesting that DiffGR can effectively detect subtle TAD differences.

### 3.2 DiffGR performed stably against changes in noise and coverage levels

Next we sought to evaluate the robustness of our method under various noise levels and sequencing coverage conditions.

In the earlier simulations, we added 10% noise to the simulated differential contact matrices. To evaluate the performance of our method under different noise levels, we fixed the proportion of altered TADs at 50% and the proportion of intra-TAD alternation at 100%, and simulated the differential contact matrices with a wide range of noise levels (1%, 5%, 10%, 20%, 50%, and 80%). Intuitively, a good detection method should easily discover the differential regions in the less noisy matrices, and it becomes more challenging to detect the differential regions in the noisier cases. Our results demonstrated that DiffGR was able to correctly rank the simulated datasets. We observed a monotonic increasing trend of the false detection rate and a decreasing tendency of other precision measures as the noise levels raised (Figure 2C and Supplementary Table S3). With moderate noise levels that were not greater than 20%, the accuracy of DiffGR remained above 0.93, indicating that our method can correctly detect differential TAD regions in such noisy cases.

The sequencing coverage of the Hi-C contact maps is another major factor that could affect the performance of our method. Considering two Hi-C replicates that have the same underlying TAD structures but different sequencing coverage levels, we questioned whether our DiffGR method can correctly categorize them as non-differential. In other words, we intended to estimate the false positive rates caused by low-coverage and sparse Hi-C data. To directly investigate the influence of the sequencing coverage on the detection of differential regions, we utilized the GM12878 chromosome 1 contact matrix as the original matrix, and generated a series of down-sampled contact matrices with lower coverage levels (20%, 40%, 60%, 80%, and 100%). Figure 2D and Supplementary Table S4 show that the average false detection rates remained below 0.05 for most coverage levels, except for the lowest coverage level of 20%, demonstrating the robustness of our DiffGR method under low-coverage conditions.

### 3.3 DiffGR successfully detected hierarchical-TAD changes

In addition to single-TAD differences, hierarchical-TAD changes also exist in some genomic regions between different cell types. In these regions, one of the Hi-C contact maps exhibits a single dominant TAD structure, while the other Hi-C contact map presents two or more subTADs separated by additional boundaries in between. Hierarchical TADs are computationally challenging to detect. Although the two Hi-C maps have different TAD boundaries, the chromatin interaction patterns within the subTADs could be very similar. Consequently, the correlation coefficients (CCs) for the strata with small genomic distances might still remain high between two contact maps. In addition, as the genomic distance increases, the weight of the corresponding stratum in the SCC calculation gradually declines. As a result, the SCC values are primarily contributed by CC values from strata with smaller genomic distances, which makes it difficult to detect differential regions in the hierarchical-TAD cases.

To evaluate the performance of DiffGR in this more challenging situation, we simulated contact matrices containing hierarchical-TAD structures with respect to varying noise levels (see Methods) and then computed the false detection rate in a similar manner as in the single-TAD simulations. As demonstrated in Figure 3 and Supplementary Table S5, the trend of the false detection rates and other measure statistics across various noise levels under the hierarchical-TAD setting was similar to the pattern observed in the single-TAD case (Figure 2C and Supplementary Table S3). Furthermore, the false detection rates remained lower than 0.05 when the noise level was within 50%. Taken together, these results indicated that DiffGR can reliably detect the differentially interacting genomic regions with hierarchical-TAD patterns.

**Figure 3:**
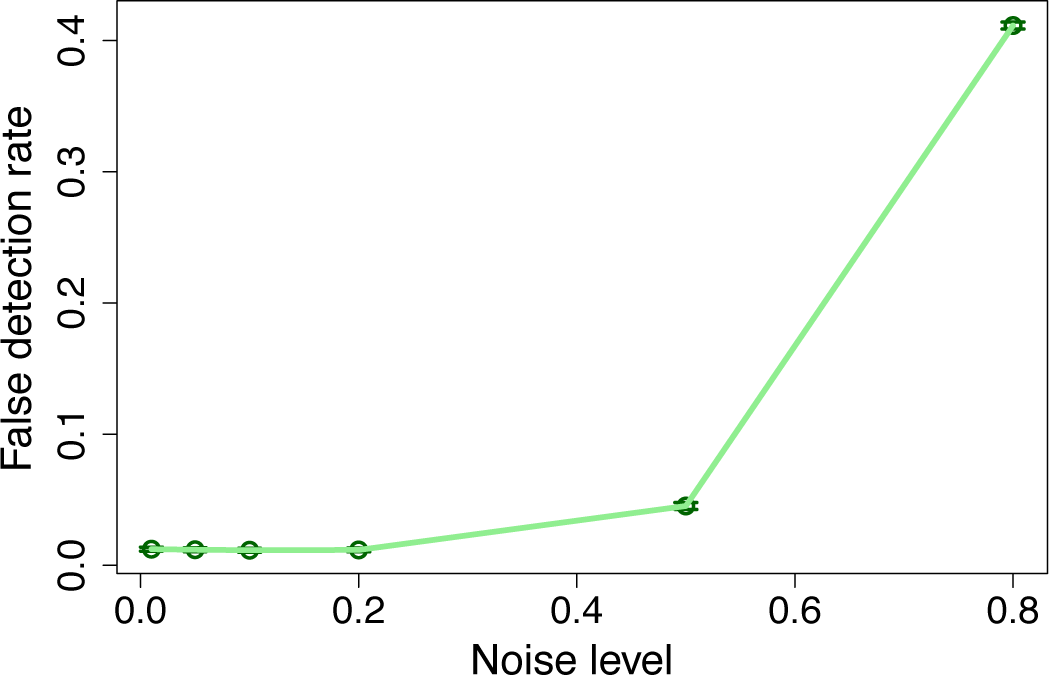
Performance of hierarchical-TAD simulations. The curve shows the mean false detection rates at various noise levels. Vertical bars represent 95% confidence intervals.

### 3.4 DiffGR revealed cell type-specific genomic interacting regions

Besides validating our method on simulated datasets, we further applied DiffGR to detect cell type-specific differences in five human cell types (GM12878, HMEC, HUVEC, K562, and NHEK) [9] and in two mouse cell types (ES and cortex cells) [11]. In total, we conducted two comparisons between biological replicates in human GM12878 and mouse ES cells, and eleven pairwise comparisons between different cell types (ten pairs among five human cell types and one pair between two mouse cell types). In each pairwise comparison, we first applied HiCseg to identify TAD boundaries from the 50-kb contact matrix for each data and then partitioned the genome into three types of candidate regions: single-TAD candidate regions, hierarchical-TAD candidate regions, and complex-TAD candidate regions. Statistically significant differential genomic regions were identified between each comparison with FDR cutoff 0.05.

We first sought to evaluate the performance of our method on biological replicates of Hi-C data. Previous studies have shown that the high degree of similarity between biological replicates and dominant consistence between TAD boundaries in replicate data [9, 11, 40]. For the comparison between human GM12878 replicates, consistent with our expectations, the majority (89.55%) of the 2325 candidate genomic across the genome regions belonged to single-TAD type and very few (2.45%) candidate genomic regions were detected as differential by our method (Supplementary Figure S2). Specifically, only 1.97% of single-TADs were identified as differential, whereas 6.17% and 4.94% were detected in hierarchical-TAD and complex-TAD cases respectively. Similar results were also witnessed in the comparison between replicates in mouse ES cells: 83.42% candidate genomic regions were classified as single-TAD type and few (6.02%) were identified as differential (Supplementary Table S6). Overall, our DiffGR results confirmed that these biological replicates displayed highly consistent chromatin structures with minor biological variations.

Next, we applied DiffGR to detect cell-type-specific differences. As illustrated in Figure 4 and Supplementary Material Table SS1, for the ten pairwise comparisons among human cell types, 55.57% of all candidate genomic regions belonged to the single-TAD category (consistent with previous observations indicating that TAD boundaries are stable across cell types [11]), 31.88% to the hierarchical-TAD category, and 12.55% to the complex-TAD category. Our DiffGR analyses showed that only 24.26% of the single-TAD candidate regions showed statistically significant differences between two samples; 59.24% of the hierarchical-TAD candidate regions were determined to be differential; while the differential proportion of the complex-TAD category was as high as 89.82%. The differential results were largely consistent when the default TAD caller was changed from HiCseg to CHDF or TADreg, demonstrating the stability of the DiffGR algorithm over different TAD callers (Supplementary Results S2). In addition, we found that the proportion of detected differential regions varied largely across chromosomes, ranging from 0.14 to 0.76 (Supplementary Figure S3). For the comparison between mouse ES and cortex cells, 20.22% of the candidate genomic regions in the single-TAD category were identified as differential, while the proportion increased to 75.94% in the complex-TAD category (Supplementary Table S7). These observations indicated that candidate genomic regions with more distinct patterns of TAD boundaries are more likely to be detected as differential between two Hi-C samples.

**Figure 4:**
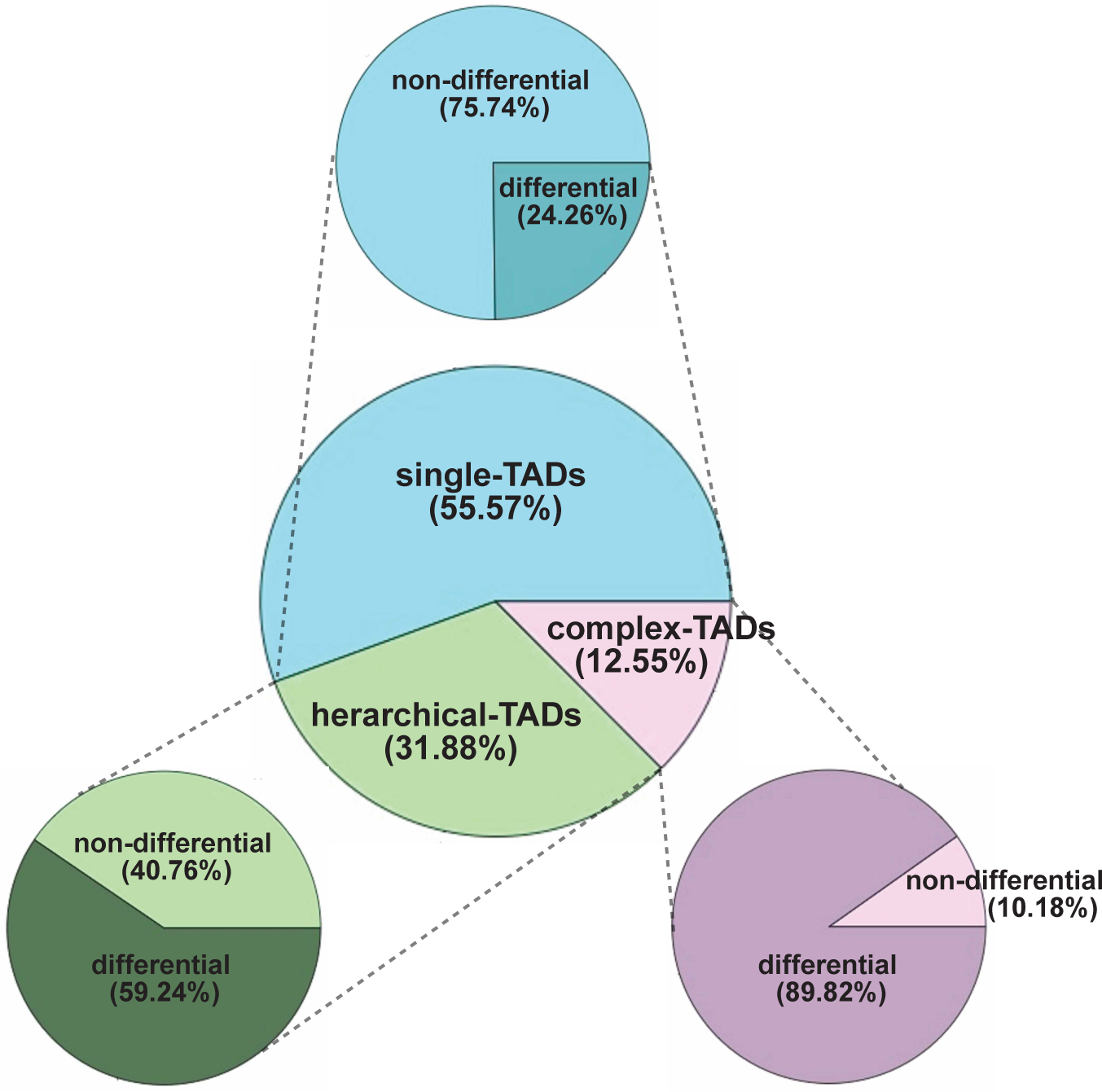
Piecharts of DiffGR results obtained from human Hi-C datasets. The DiffGR results from the ten pairwise comparisons among five human cell types (GM12878, HMEC, HUVEC, K562, and NHEK) [9] using the TAD caller HiCseg are summarized. The center piechart presents the proportions of three categories of candidate regions. The three outer piecharts display the proportions of DiffGR-detected differential genomic regions, one for each candidate category.

In addition to partitioning the genome at 50-kb resolution, we also performed differential analyses on the five human Hi-C datasets at 25-kb and 100-kb resolutions, separately. We calculated the overlapping rate (that is, the proportion of the genome that was classified into the same differential or non-differential status) between different resolutions. Overall, we observed a high consistency between the detected differential regions across different resolutions, where the overlapping rate was 0.8207 between the detection results at 50-kb and 100-kb resolutions, 0.8956 between those at 25-kb and 50-kb resolutions, and 0.7712 between those at 25-kb and 100-kb resolutions. These results demonstrated that DiffGR can robustly and consistently detect cell type-specific differential genomic regions across various resolutions.

### 3.5 Changes in CTCF and histone modification patterns were consistent with DiffGR detection results

As there is no ground truth of differential chromatin interacting regions in real data, we sought to evaluate the performance of our method by investigating the association between the changes in 1D epigenomic features and 3D genomic interaction regions. The chromatin architectural protein CTCF plays an essential role in establishing higher-order chromatin structures such as TADs. In addition, it has been shown that transcription factors and histone marks are enriched or depleted at TAD boundaries and are associated with active enhancers, promoters and transcribed genes [11, 12]. Therefore, we expected that differential bindings of transcription factors such as CTCF and histone modifications would be more likely to located in differential genomic interacting regions.

To test this hypothesis, we then utilized the ChIP-seq datasets of CTCF and histone modifications from the ENCODE project [41]. For each ChIP-seq dataset, we called the peaks via MACS2 [43] and detected differential peak by DEseq2 [25]. Then, we calculated the proportions of differential peaks that were located in DiffGR-detected differential genomic regions at 100-kb, 50-kb, and 25-kb resolutions. Further, we checked the significance of differential peak enrichment by randomly selecting a bundle of peaks (where the peak number is the same as the number of differential peaks detected by DEseq2) with 2000 times and calculating their corresponding percentages located in differential genomic regions to estimate the P-values.

Table 1 summarizes the ChIP-seq analyses on the DiffGR detection results obtained from five human Hi-C datasets [9]. Overall, DiffGR-detected differential genomic regions were supported by 1D epigenomic features. In particular, we observed that the agreement between the changes in ChIP-seq signal and chromatin structures was improved in finer-resolution analyses. As shown in Table 1, 52.48% of the differential CTCF peaks appeared in DiffGR-detected differential genomic regions at 100-kb resolution; whereas in the results at 25-kb resolution, 74.85% of differential CTCF peaks were located in differential genomic regions. In addition, the histone modification datasets showed similar results concordant with the detection results of differentially interacting regions in Hi-C contact maps. At 25-kb resolution, the majority (*>* 70%) of differential histone peaks showed significant consistency with differentially interacting regions for all four histone modification datasets, including H3K4me2, H3K9me3, H3K27ac, and H3K27me3. Collectively, these results indicated that the changes in CTCF bindings and histone modifications were in good agreement with the differences in genomic interacting regions. Furthermore, at finer resolution DiffGR produced more accurate identification of differentially interacting genomic regions in higher agreement with the CTCF and histone modification data.

**Table 1:**
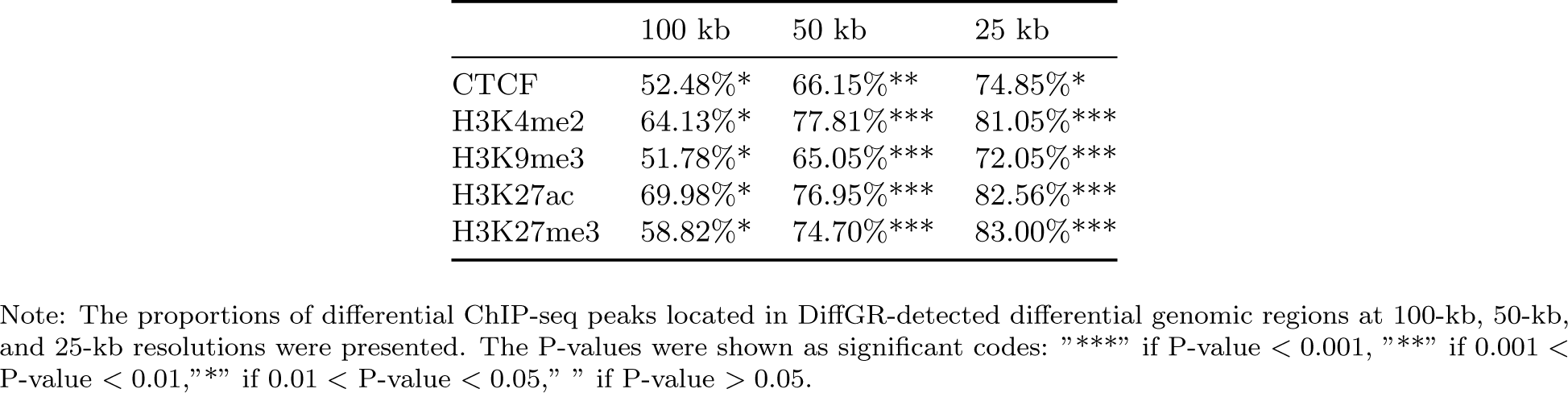
Agreements between ChIP-seq data and DiffGR-detected differential genomic regions in human Hi-C datasets.

### 3.6 Differential RNA-seq analysis results were consistent with DiffGR detection

In addition to investigating the changes in 1D epigenomic features, we further studied the relationship between quantitative changes in gene expression levels and 3D genomic interaction regions. Previous studies have showed that topological changes of 3D genome organization have a large effect on the cross-talk between enhancers and promoters therefore can alter gene expression [9, 22]. Thus, we expected to observe an enrichment of differential expressed genes in DiffGR-detected differential genomic regions.

To evaluate this assumption, we first detected significant changes in gene expression levels between human GM12878 and K562 cells using DESeq2 [25] and those between mouse ES and cortex cells using ballgown [44]. Then we calculated the percentage of differentially expressed genes that were located inside the DiffGR-identified differential genomic regions. To calculate the enrichment of differentially expressed genes, we randomly chose a set of genes, whose number is equivalent to the number of the DESeq2-detected differentially expressed genes, with 200 times, computed their corresponding proportions located in differential genomic regions, and then performed *t*-test for comparison. In summary, a total number of 8781 differentially expressed genes were detected between human GM12878 and K562 cells, and 79.54% of them were located in DiffGR-detected differential genomic regions (*p*-value = 3.72 *×* 10^−5^, permutation test); whereas 2124 differentially expressed genes were identified between mouse ES and cortex cells and 61.66% were within DiffGR-detected differential genomic regions (*p*-value *<* 2.2 *×* 10^−16^). Taken together, these results demonstrated that the changes of gene expression in RNA-seq data were highly consistent with the DiffGR detection results.

To further explore the potential functional roles of the genes located in differential genomic regions detected by DiffGR, we performed Gene Ontology (GO) enrichment analysis on those genes using DAVID [45]. As shown in Table 2, we observed a high enrichment of GO terms related to the immune responses, which is consistent with the immunological nature of GM12878 lymphoblastoid B-cells.

**Table 2:** Functional enrichment of genes located in differential genomic regions between GM12878 and K562.

| GO Term | FDR |
| --- | --- |
| GO:0046649 lymphocyte activation | 2.7E-11 |
| GO:0002376 immune system process | 4.6E-11 |
| GO:0002520 immune system development | 2.3E-9 |
| GO:0070663 regulation of leukocyte proliferation | 5.2E-7 |
| GO:0042113 B cell activation | 7.7E-7 |
| GO:0030183 B cell differentiation | 2.2E-5 |

Note: Genes located within differential genomic regions at 25-kb resolution were utilized in GO enrichment analysis. Test results were reported by DAVID [45].

### 3.7 DiffGR detection was supported by differential chromatin interactions

Several Hi-C comparative studies have demonstrated that the majority of the chromatin structural changes tend to couple with the formation/disappearance of topologically associated domains (TADs) [9, 22], implying that changes in Hi-C interaction counts are likely to be observed within genomic regions at TAD level. Hence, we checked differential chromatin interactions (DCIs) between GM12878 and K562 cells at 50-kb resolution by FIND [28] and compared FIND results with our DiffGR results. As shown in Figure 5, the percentages of DCIs detected by FIND located within candidate genomic regions were dominant in the majority of chromosomes and with 55.43% across the whole genome. In addition, 82.80% of the DCIs located in candidate genomic regions are classified into differential regions, demonstrating that DiffGR effectively detected the regions with significant changes in chromatin contacts.

**Figure 5:**
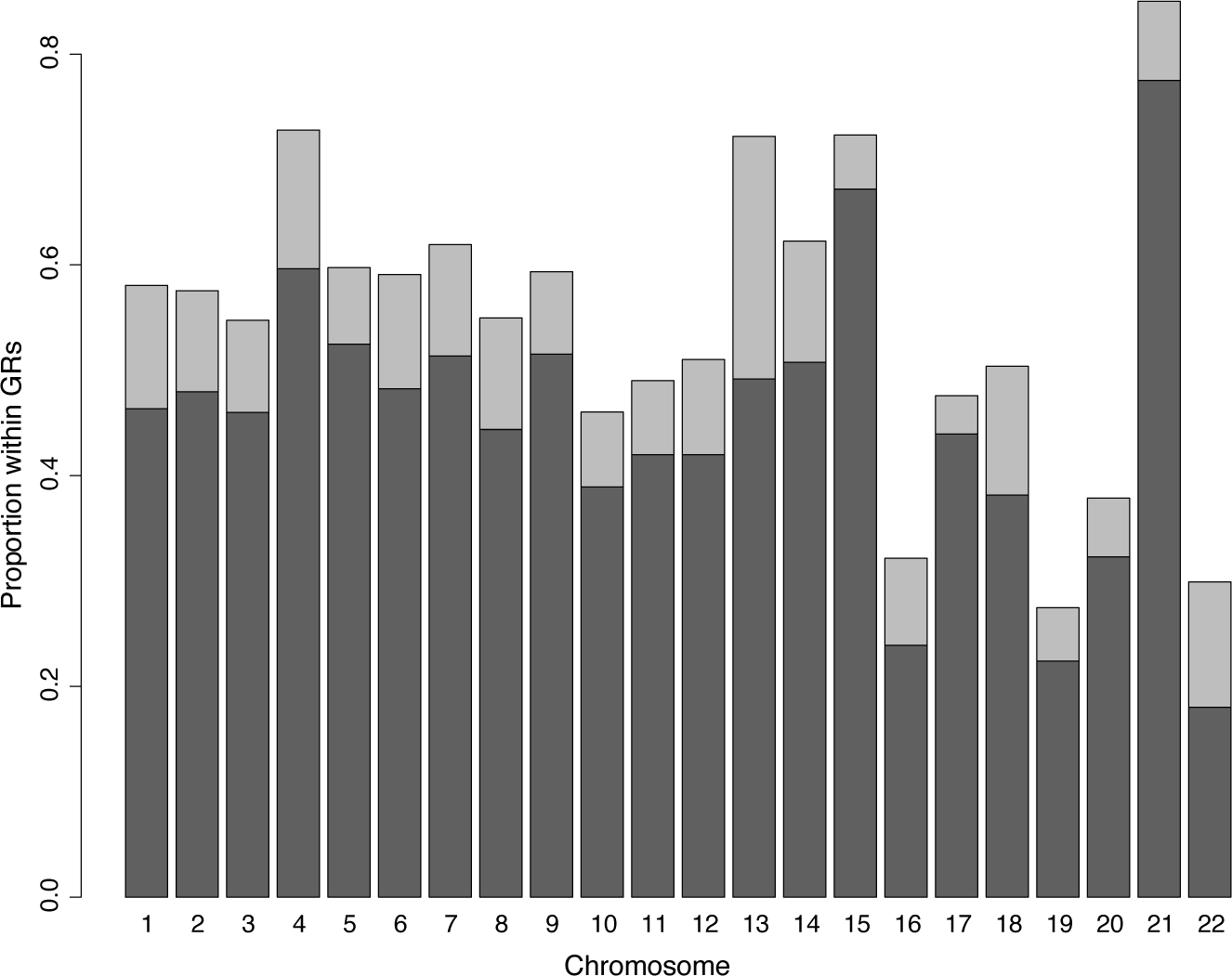
Comparison between FIND and DiffGR. Barchart of the proportions of FIND-detected DCIs located in candidate genomic regions and differential genomic regions for all autosomes between GM12878 and K562. The light gray bars denote the proportions of DCIs located in candidate genomic regions; the dark gray bars represent the proportions of DCIs located in differential genomic regions.

### 3.8 Performance and comparison with state-of-the-art differential TAD detection tools

To further investigate the performance of DiffGR, we compared the DiffGR results with three differential TAD boundaries detection methods (HiCDB [13], TADCompare [31], and TADreg [32]) on both simulated and real data. In the simulation part, we compared all four methods using the synthetic data under the default setting (proportion of altered TADs = 50%, proportion of TAD alternation = 100%, and noise level = 10%) and calculated corresponding sensitivities and specificities. As shown in Table 3, the sensitivities of the three differential TAD boundaries detection methods are all above 70% and comparable to the sensitivity of DiffGR, while the specificities of the three differential TAD boundaries detection methods are relatively low. These results demonstrated that HiCDB, TADCompare, and TADreg can accurately identify most differential TAD boundaries within differential regions, but falsely detect many non-differential boundaries within differential regions. We would like to point out that the simulation process were designed to check the robustness of DiffGR detection results by generating random structures within the predefined differential areas and keeping the original chromatin structures within non-differential regions. Therefore, differential TAD boundaries are expected to appear within differential regions, while non-differential TAD boundaries may located either within or outside the non-differential regions. As a result, the performance on specificities of the differential TAD boundary detection tools were affected.

**Table 3:** Simulated Data Comparison among differential TAD boundaries tools and DiffGR.

|  | Sensitivity |  | Specificity |  |
| --- | --- | --- | --- | --- |
|  | Mean | SD | Mean | SD |
| DiffGR | 0.8871 | 0.0239 | 0.9904 | 0.0030 |
| HiCDB | 0.7758 | 0.0193 | 0.0630 | 0.0415 |
| TADCompare | 0.9568 | 0.0113 | 0.3128 | 0.0284 |
| TADreg | 0.7762 | 0.0660 | 0.3124 | 0.0303 |

Next, we compared the DiffGR results with three differential TAD boundaries detection methods on the five human Hi-C datasets by Rao et al. [9] and the two mouse datasets by Dixon et al. [11] at 50-kb resolution. Overall, the differential TAD boundaries identified by HiCDB, TADCompare, and TADreg were highly concordant with DiffGR-detected differentially interacting genomic regions. Notably, 73.86% of the HiCDB-detected, 76.25% of the TADCompare-detected, and 71.90% of the TADreg-detected differential TAD boundaries displayed consistent results with our DiffGR detection in the human datasets. In addition, highly concordant rates were also witnessed in the mouse dataset with 59.56%, 62.01%, and 60.32% consistency rate with HiCDB, TADCompare, and TADreg, respectively.

Furthermore, we compared DiffGR with HiCExplorer [33–35]), the only available tool for differential TAD regions detection, on the five human Hi-C datasets by Rao et al. [9] at 50-kb resolution. We observed that 60.62% of the 2877 HiCExplorer-identified differential regions overlapped with DiffGR-detected differential regions. To better compare the detection results of DiffGR and HiCExplorer, we then computed the concordant rates between HiCExplorer-detected differential regions and differential ChIP-seq peaks. As shown in Table 4, in comparison with DiffGR results, we observed a higher proportion of differential ChIP-seq peaks located in HiCExplorer-detected differential regions but most of the results are not statistically significant, indicating that HiC-Explorer identified a great amount of differential regions, however partial detected regions are not significantly differential to some extents. Further, we investigated the advantages of DiffGR and HiCexplorer over TADCompare results, and found that differential TAD boundaries within DiffGR-detected differential regions have a larger proportion located closer to the CTCF and other histone differential peaks than those outside differential areas, while some disagreements were found in HiCexplorer-detected differential regions(Supplementary Results S3). Collectively, these results indicated that DiffGR-detected differential genomic regions had a better agreement with 1D epigenomic features than HiCExplorer-detected differential regions.

**Table 4:** Agreements between ChIP-seq data and differential genomic regions detected by DiffGR and HiCExplorer.

|  | DiffGR | HiCEXplorer |
| --- | --- | --- |
| CTCF | 66.15%** | 86.48% |
| H3K4me2 | 77.81%*** | 86.11% |
| H3K9me3 | 65.05%*** | 86.96%*** |
| H3K27ac | 76.95%*** | 86.10% |
| H3K27me3 | 74.70%*** | 85.21% |

Note: The proportions of differential ChIP-seq peaks located in differential genomic regions detected by DiffGR and HiCEXplorer at 50-kb resolution were presented. The P-values were shown as significant codes: "\*\*\*\*" if P-value < 0.001, "\*\*\*" if 0.001 < P-value < 0.01, "\*\*" if 0.01 < P-value < 0.05, " " if P-value > 0.05.

### 3.9 Role of chromatin structure in K562 differentiation

MYC binding protein2 (MYCBP2) encodes an E3 ubiquitin-protein ligase and reduced expression of this gene was observed in leukemia patients, which has been revealed that CK2 inhibitor take the anti-leukemia effect through Ikaros-mediated regulation on MYCBP2 expression in high-risk leukemia[46].

In Figure 6, we marked the differential genomic region reported by DiffGR, which is a hierarchical TAD region showing a large TAD in K562 cell and two sub-TADs in GM12878 cell. We notice that the interacting region located around the unique TAD boundary of GM12878 show a significant differential binding of the CTCF protein given by DESeq2. RNA-seq tracks at this region also indicate an obvious loss of expression in K562 cell (immortalised myelogenous leukemia cell line) compared to GM12878 cell(B-Lymphocyte cell), demonstrating MYCBP2 low expression is correlated with acute leukemia.

**Figure 6:**
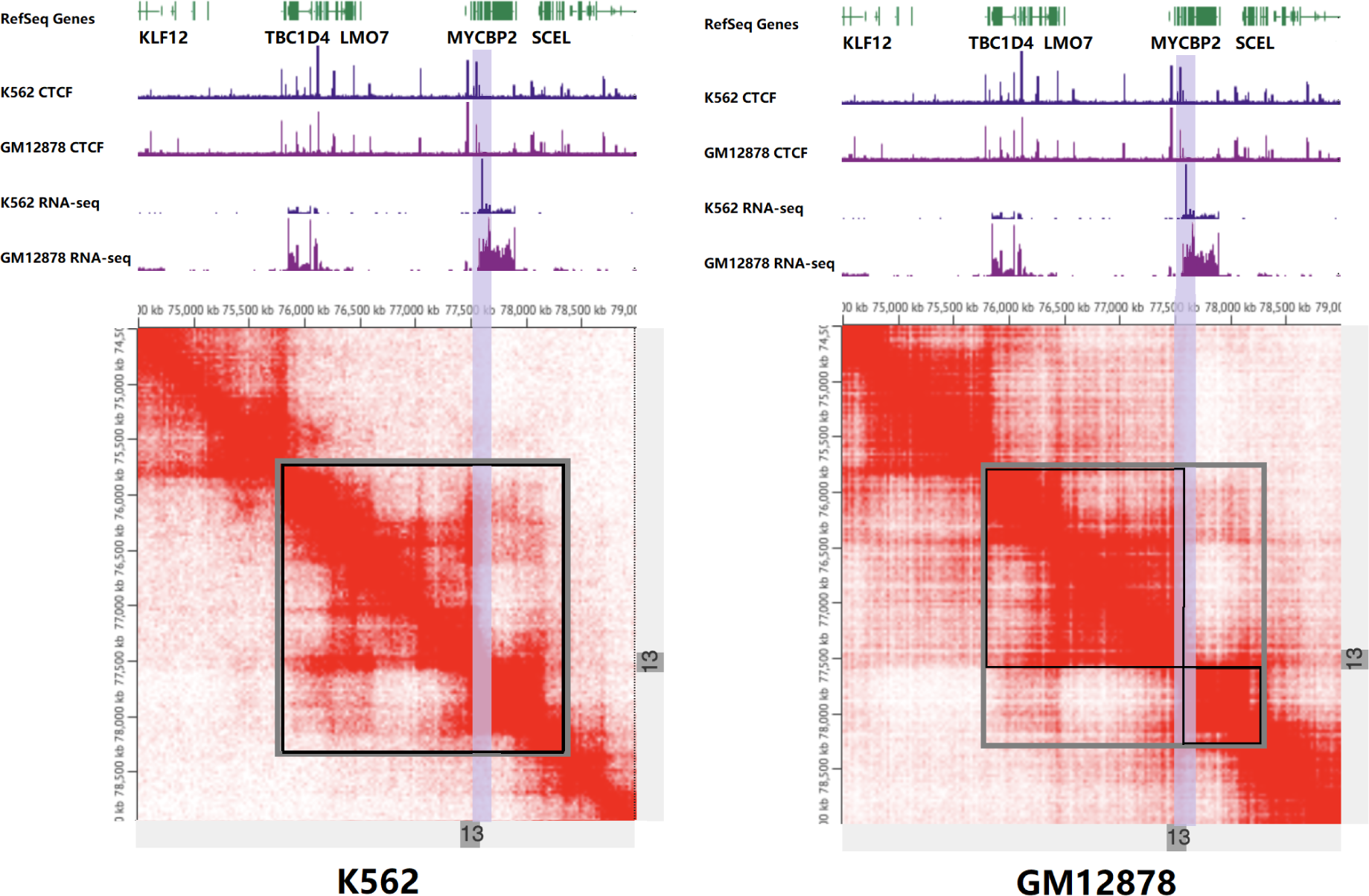
Concrete example of differential regions between GM12878 and K562. Hi-C contact map of the K562 and GM12878 cells at chr13: 74,500kb-79,000kb are displayed. Differential genomic region (with SCC value 0.6551 and pvalue 0.0334) is shown in gray square and the TADs for each cell types are shown in black squares. The differential CTCF peak region is highlighted by the purple bars.

## 4 Discussion and Conclusions

With the fast accumulation of Hi-C datasets, there has been a dramatically increasing interest in comparative analysis of Hi-C contact maps. However, most existing methods for comparative Hi-C analysis focused on the identification of differential chromatin interactions, while few studies addressed the detection of differential chromatin organization at TAD scale. To tackle this problem, we developed a novel method, DiffGR, for calling differentially interacting genomic regions between two Hi-C contact maps. Taking genomic distance features of Hi-C data into consideration, our algorithm utilized the SCC metric instead of the standard Pearson CC to measure the similarity of local genomic regions between Hi-C contact maps. Furthermore, we proposed a nonparametric permutation test to assess the statistical significance of the local SCC values. In contrast to the parametric approaches that were used by most Hi-C data analysis methods, our nonparametric approach does not have a set of predefined assumptions about the nature of the null distribution and, therefore, is more robust and can be applied to more diverse data from real cases. Additionally, we utilized a non-parametric smoothing spline regression to speed up the permutation test and showed that the speed-up algorithm can steadily produce consistent outputs. Through empirical evaluations, we have demonstrated that DiffGR can effectively discover differential regions in both simulated data and real Hi-C data from different cell types. That is, DiffGR produced robust and stable detection results under various noise and coverage levels in simulated data; DiffGR detection results in real data were effectively validated by the ChIP-seq and RNA-seq data; DiffGR produced consistent and advantageous results compared with state-of-the-art differential TAD boundaries/regions detection tools. To summarize, DiffGR provides a statistically rigorous method for the detection of differentially interacting genomic regions in Hi-C contact maps from different cells and conditions, therefore would facilitate the investigation of their biological functions.

We envision a few possible extensions and future directions based on this work. First, our method performs pairwise comparison between Hi-C contact maps. One potential future direction is to design a more general statistical framework for differential analyses among three or more samples. Then we could further assign the differentially interacting genomic regions to cell type-specific or condition-specific changing areas. Second, we currently pool biological replicates together in our analyses. Extending DiffGR to incorporate multiple biological replicates to detect reproducible differences would enhance the reliability of the detection results. Third, in our algorithm, we use the shared TAD boundaries between two samples to segment the genome into candidate genomic regions and then detect differential regions. Recently, the notion of TADs being highly conserved across cell types has been questioned [47, 48]. Therefore, a more general approach to define and classify the candidate genomic regions would be beneficial to better characterize the variability of chromatin interactions between different conditions. Lastly, our method is specifically designed for bulk Hi-C data. Given the high sparsity and variability of single-cell Hi-C contact matrices, identifying differential genomic regions at single-cell level remains a significant challenge.

## 5 Code Availability

The DiffGR R Code (both algorithm and simulation) is publicly available at https://github.com/ wmalab/DiffGR under the GNU GPL ≥ 2 license. The source code is also available at BioCode https://ngdc.cncb.ac.cn/biocode/tools/BT007313.

## 6 CRediT author statement

**Huiling Liu**: Conceptualization, Methodology, Software, Formal analysis, Writing - Original Draft. **Wenxiu Ma**: Conceptualization, Supervision, Writing - Review & Editing, Funding acquisition. All authors read and approved the final manuscript.

## 8 Acknowledgments

The authors would like to thank Tiantian Ye, Yangyang Hu, and Luke Klein for helpful discussions and feedback, and the editor and reviewers for their insightful comments and suggestions. This work was supported by the National Science Foundation, USA [DBI-1751317]; and the National Institute of Health, USA [R35GM133678].

